# Secondary sexual dimorphism traits under abiotic stress in dioecious species: The case of *Amaranthus palmeri*

**DOI:** 10.1101/2022.02.15.480572

**Authors:** Nicholas E. Korres, Jason K. Norsworthy, Toby FitzSimons, Trenton L. Roberts, Derrick M. Oosterhuis, Govindjee Govindjee

**Affiliations:** Dept. of Agriculture, University of Ioannina, Kostakii, Arta 47100, Greece; Crop Soil and Environmental Sciences, University of Arkansas, Fayetteville, AR 72704, USA; PepsiCo Inc., St. Paul, MN 55108, USA; Biochemistry, Biophysics and Plant Biology, University of Illinois at Urbana-Champaign, Urbana, IL 61801, USA

**Keywords:** Chlorophyll, Chlorophyll fluorescence, Gender, Light intensity, Mineral deficiency, PS II capacity, Photosynthetic efficiency

## Abstract

The evolution of secondary sex-specific traits of dioecious species under abiotic stress conditions has received limited research, especially in the case of *Amaranthus palmeri*, a fast adapting and highly competing plant. Here, we have examined the interactive effects of abiotic stress on mineral accumulation, chlorophyll *a* and *b* content, and the operating capacity of Photosystem II (PSII) in both male and female *A. palmeri* plants grown under three different intensities (150, 450 and 1300 μmol photons m^−2^ s^−1^) of white light, and under N, K or P deficiency. Mineral profiling of the leaves and stems (with inflorescence) highlighted intra- and intersexual differences in their accumulation pattern and mineral associations. Chlorophyll *a* and *b* content was different between the male and the female plants, being lower in the latter, at high light intensity, especially as the flowering progressed, or when they were under K or P deficient condition. Further, the chlorophyll *a*/*b* ratio was lower at the higher light intensity in the female, over that in the male, plants. Chlorophyll fluorescence parameters, i.e., steady state (F’_S_) and maximum (F’_M_) fluorescence increased under high light intensity, whereas the PSII operating efficiency (Φ_PSII_) decreased in the female plants, indicating reduced PSII capacity. Sex-specific differences in *A. palmeri* showed a differential response to stressful conditions because of differences in their ontogeny and physiology, and possibly due to the cost of reproduction. We suggest that the breeding system of dioecious species has weaknesses that can be used for the ecological management of dioecious weed species.

## Introduction

*Amaranthus palmeri*, one of the most problematic weeds in many row and vegetable crops worldwide, exhibits remarkable biological and physiological characteristics such as rapid growth rate, high prolific capacity, and great adaptability under biotic (e.g., crop competition) or abiotic (e.g., environmental, or chemical) stressful conditions (Webster and Nicholas 2021; Mesgaran et al., 2021). These characteristics enhance its invasive potential, competitive ability (Webster and Nichols, 2012; Korres et al. 2019; Korres et al., 2020) and its capability to develop multiple herbicide resistance to a range of herbicide families (Heap, 2021) resulting in high infestation levels of natural habitats and agroecosystems (Korres et al., 2019; Korres et al., 2020). Consequently, its presence leads to food production cost increases (Korres et al., 2019).

As a summer ephemeral species, *A*. *palmeri* grows well under xerophytic and heliophytic conditions (Korres et al., 2017). It prefers warm or moderate temperatures and thrives in soils rich in nitrogen content (Korres et al., 2017). However, climate change imposes a strong selection pressure on weeds (Ziska et al., 2019) that eventually adapt rapidly to changes through alterations in their biology, physiology, and phenology (Korres and Dayan, 2020). This is particularly true for C4 weed species, as is the case of *A. palmeri*, which has already expanded its habitable range in the northern states of USA, as well as in the southern Canadian provinces (Korres and Dayan, 2020). Fitzpatrick et al. (2017) have emphasized the need for trait-based studies since phenotypic variations can affect plant-environment trade-offs, thus deepening our understanding of species interaction with its environment and consequently the species habituation. However, the habituation of a species, an important phenomenon *per se*, does not intrinsically expand the behavioral repertoire of an organism (Adelman 2018). This is of a particular importance since the obligatory outcrossing of *A. palmeri*, due to its dioecious nature, results in high genetic diversity, broad adaptability, and enhanced evolutionary capacity (Franssen et al., 2001; Montgomery et al., 2021). Furthermore, *A. palmeri* is known to exhibit great plasticity to external stimuli and has distinguishable sex differentiation under a wide range of environments (Korres and Norsworthy, 2017; Korres et al., 2017). Research on the plasticity of plants has revealed morphological and allocation responses to environmental stimuli along with ontogenetic trajectories and offspring traits (Sultan, 2005). Further, dioecious species have evolved sex-specific secondary traits, i.e., secondary sexual dimorphism and functional differences (Dudley, 2006), some of them attributed to differences in their reproductive functions (Obeso, 2002), but also to physiological and ontogenetic functions during their vegetative growth (Sanchez-Vilas and Retuerto, 2012; Zhang et al., 2011; Montesinos et al., 2012). For example, greater height and biomass production in the female than in the male *A. palmeri* plants grown under field conditions have been reported (Korres et al., 2020; Webster and Gray, 2015) (Supplement Material-SM-, Fig. S1).

This differential response is a fitness strategy by the female plants in their “effort” to reproduce (Obedso, 2002, Solbrig, 1981) and related to physiological functions of the plant such as photosynthetic performance, water use efficiency or even phenology (Rowland and Johnson, 2001; Delph, 1999). Another characteristic of dioecious species, “division of labor” (Charnov, 2016), is expressed through the manifestation of resource acquisition and allocation. In complex and ever-changing environment, resources are often scarce with unequal spatial and temporal distribution (Gagliano et al., 2016). As plants are simultaneously exposed to diverse abiotic stresses, such as mineral deficiency, and/or inadequate light environment, they have developed specific mechanisms to tackle different types of abiotic stress they are exposed to during their growth (Ramegowdaa and Senthill-Kumar, 2005).

Deficiency of any of the essential minerals such as nitrogen (N), phosphorous (P) and potassium (K) restricts plant growth and development through reduction in photosynthetic efficiency. However, plants are concurrently exposed to various abiotic stresses such as mineral deficiency and varying regimes of light intensity (Reddy, 2006). In *A. palmeri*, light environment affects acclimation through adjustments in plant ontogeny, especially of morphological traits during both its vegetative and reproductive stages (Brainard et al., 2005). Investigating and characterizing the complex interactions between weeds and their environment in natural habitats and agroecosystems is vital for understanding the dynamics of species occurrence in both natural and agricultural-based habitats (Brainard et al., 2005). To the best of our knowledge, limited research has demonstrated the differential response of dioecious male and female *A. palmeri* plants to abiotic stress related to N, P and K deficient environment, as well as to varying light intensities (Korres et al., 2017). The research described here clearly reveals key differences in sex-specific secondary traits related to biological and functional characteristics between the male and the female *A. palmeri* plants grown under different light intensities and NPK deficient environment. We suggest that the male and the female plants grown under stress conditions initiate secondary sex-specific traits related to mineral content accumulation, chlorophyll content and photosynthetic capacity. Specific research questions addressed here include: (**i**) Does *A. palmeri* mineral content accumulation differ between male and female plant organs (leaves vs. stems incl. inflorescences) when they are exposed to abiotic stress? (**ii**) Do male and female *A. palmeri* plants exhibit differences in chlorophyll content when grown under abiotic stress? (**iii**) Is the operating efficiency of PSII, as evaluated by measurements of chlorophyl fluorescence, of *A. palmeri* male plants less functionally plastic than that of the female plants under abiotic stress? Answers to these questions have revealed important information on the ontogenetic and physiological secondary traits that allow for greater competitive ability and invasive potential of dioecious species such as *A. palmeri*. However, the dioecious nature of the species conceals weaknesses that can be used for the ecological management of dioecious weeds without relying solely on the use of herbicides.

## Results

We present below experimental results related to the similarities and differences between the male and female plants of *A. palmeri* grown under abiotic stress; effects where sex is not involved are not included here.

### Inter- and intrasexual accumulation of minerals in *A. palmeri*

To begin with, analysis of the mineral content of untreated male and female *A. palmeri* plants showed no significant differences (P=0.451), both in the leaves and the stems. However, mineral content between the leaves and the stems of treated plants was significantly (P<0.001) affected by NPK deficiency, which was different in male and female *A. palmeri* plants (Table 1 and SM Table S1). Light intensity did not cause any effect on the mineral content of stems and leaves, either in the male or the female plant (SM Table S1). Under NPK deficient condition, the mineral content of male and female plants was significantly different for all the elements measured, except for Fe, Cu and B (Table 1 and SM Table S1). Mineral content in the stems, mostly under K and P deficiency, was found to be greater in the female compared to the male plants. For example, the stem N content was 46% and 91% greater in the female than that in the male plants, under K and P deficiency respectively. Similarly, Ca content in the stems of the female plants was 30% greater than that in the male plants for both K and P deficiency. Likewise, Mg was higher by 35% in the female plants compared to the male plants for each NPK treatment. Similarly higher differences were recorded for all other stem mineral contents in the female compared to the male plants except Fe, Cu and B (Table 1).

**Table 1.**
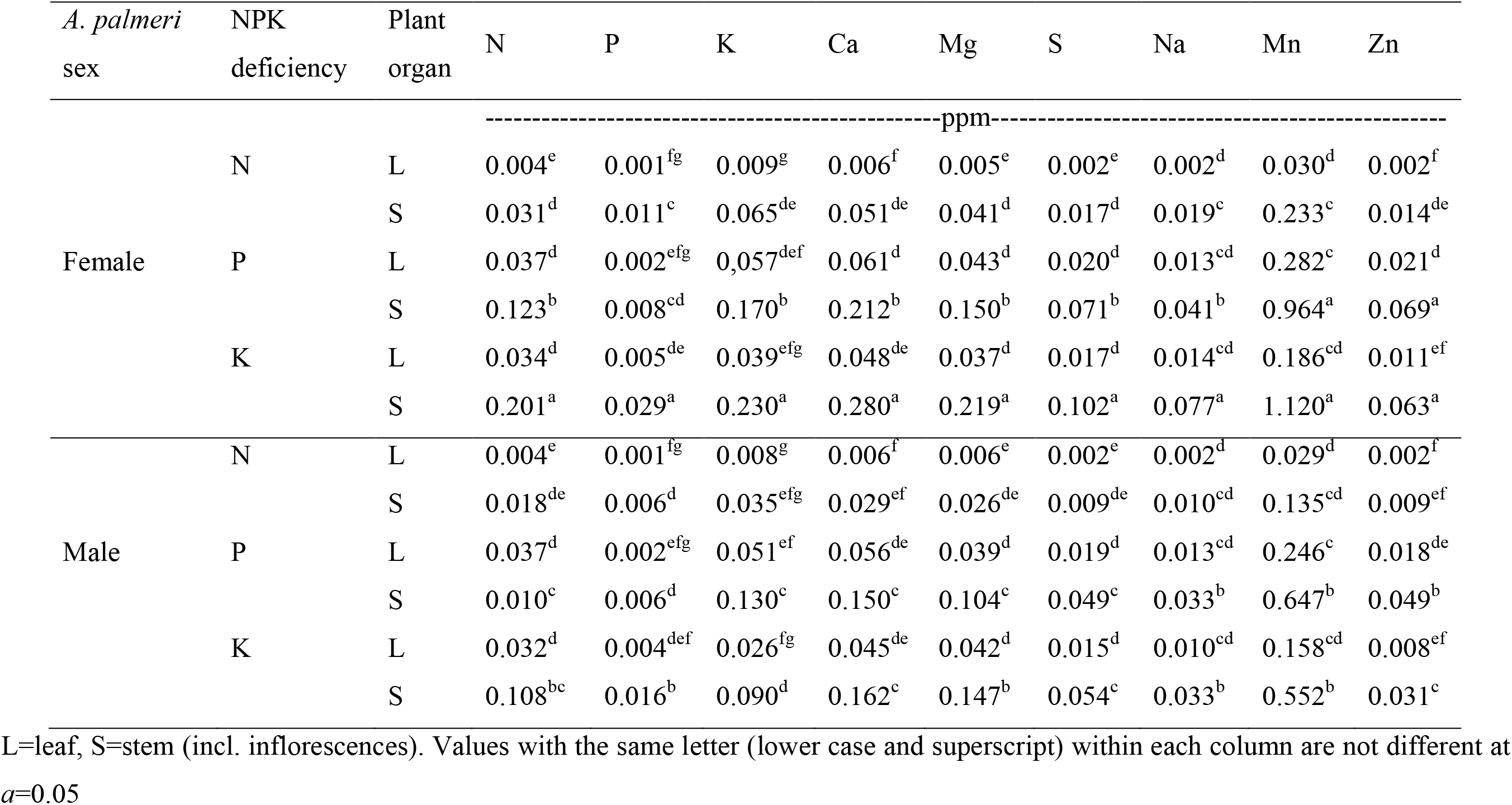
Effects of *A. palmeri* sex × NPK deficiency × plant organ on leaf and stem mineral content at harvest averaged across light intensities

Further, we observed intrasexual differences (P<0.001) between the stem and the leaf mineral content of *A. palmeri* female plants. In particular, the mineral content in the stems of the female plants was on the average 8.4-, 3.3- and 5.7-fold greater than that in the leaves under N, P and K deficiency respectively (Table 1). On the contrary, the mineral content in the stem of the male plants was 4.5-fold, 2.4-fold, and 3-fold than that in the leaves under N, P and K deficient environment (Table 1 and SM Table S1) respectively.

In the female plants, we observed positive correlations between Cu-Zn, Zn-Mn, and Cu-Fe (in the leaves), and Zn-Ca, B-Mg (in the stems) (Fig. 1A, C and SM Table S2, S3) whereas in the male plants the positive correlations were between N-P, N-K, P-Mg, Ca-S, and B-Mn contents in both stem and leaves (Fig. 1 B, D). Negative correlations between B-Fe (in stems and leaves), K- Fe (in stems only) and P-Cu (in leaves only) contents were observed in the male plants (Fig. 1 B, D and SM Tables S2, S3). In the female plants, however, negative correlations were recorded between B-Cu, Cu-Mn, Zn-S, Zn-Fe, N-S (in leaves only) and P-Zn, Zn-S and Ca-Fe (in stems only) (Fig. 1 A, C and SM Tables S2, S3). On the other hand, a negative correlation between the leaf P-K content, and positive correlations between the Mn-Fe content (in stems and leaves), N- Cu content (in leaves only) and Cu-Zn, Ca-S content (in stems only) were recorded in both male and female *A. palmeri* plants (Fig. 1 and SM Tables S2, S3).

**Figure 1.**
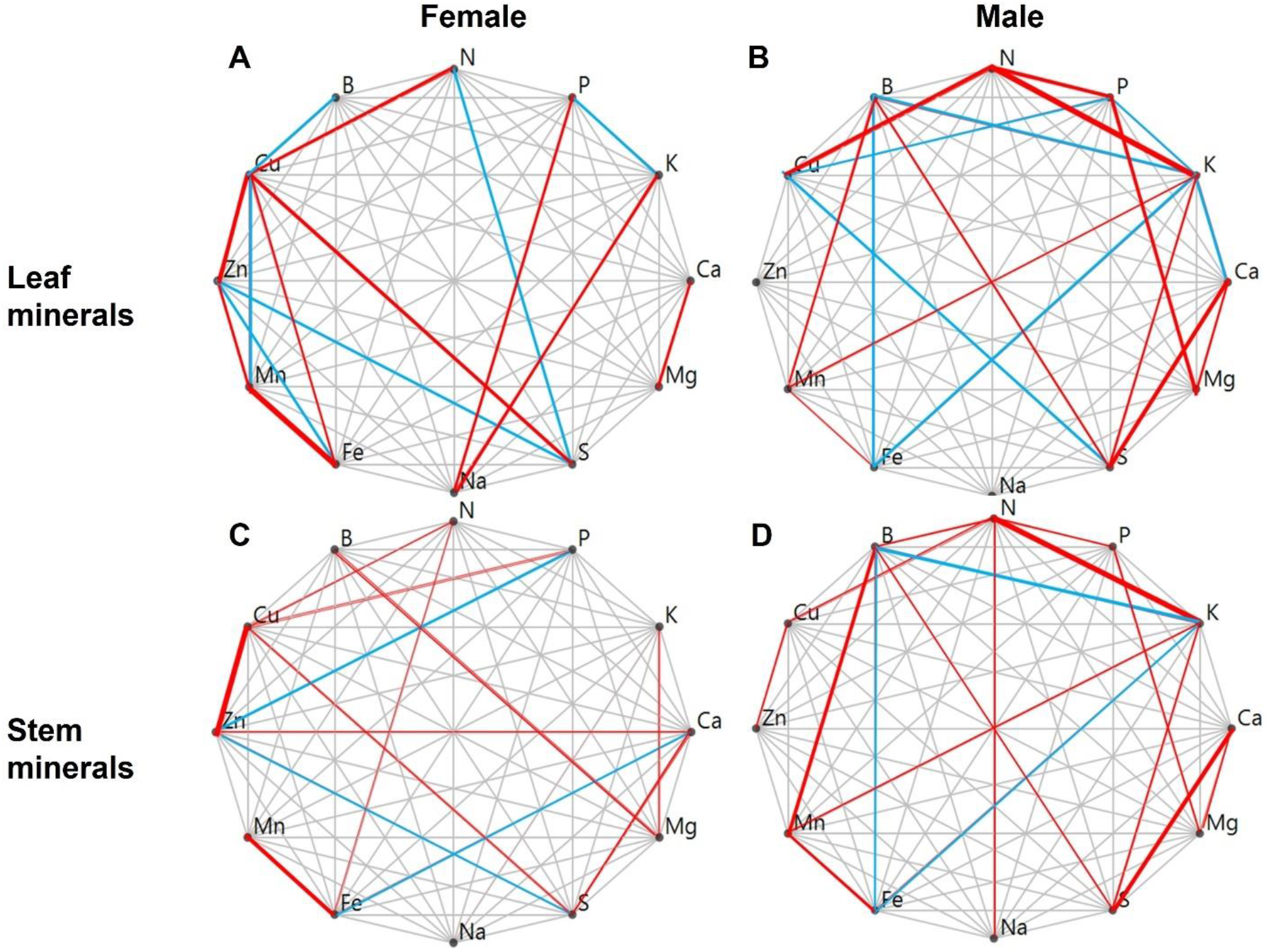
Mineral content of the leaves and the stems incl. inflorescences of the male and female *Amaranthus palmeri* plants. Red and blue lines connecting two minerals within each graph represent a positive or a negative partial correlation between these two minerals respectively. The thicker the line the higher the significance level (see Tables S2, and S3 for correlation coefficients).

In general, correlations between the micro-nutrients (i.e., Cu, Zn, Mn, S and Fe) were mostly observed in the leaves of female plants compared to that in the male plants (Fig. 1 A, B). We note that the correlations between the leaf minerals were mostly observed for the macro-nutrients (i.e., N, P, K) and the cations such as Ca, and Mg (Fig. 1 and SM Tables S2, S3). Interestingly, comparable positive and/or negative correlations between the mineral contents in the stems and the leaves, such as for N-P, N-K, P-Mg, Ca-S, B-Mn, Mn-Fe, Fe-B, were observed in male plants only (Fig. 1 B, D and SM Table S2, S). In contrast, correlations between the leaf and the stem mineral content in the female plants did not show a clear pattern for most cases, with the exception for Cu-Zn and for Mn-Fe (Fig. 1 A, C and SM Tables S2, S3).

A negative correlation between P-K was observed in the leaves, but not in the stems, of *A. palmeri* female plants. Further, the positive correlation, observed between Cu-K, Zn-Ca and Cu- P content in the stems, was not found in the leaves of female plants (Fig. 1 A, C and SM Tables S2, S3).

### Chlorophyll α and chlorophyll *b* content of male and female *Amaranthus palmeri* plants under abiotic stress

Nitrogen deficiency, when white light intensity was increased, caused a significant decrease in both chlorophyll *a* (P = 0.007) and chlorophyll *b* (P= 0006) content, averaged across the sampling times; this result was independent of the sex of the *A. palmeri* plants (Fig. 2 A, B and SM Table S4). However, as compared to the male plants, the female plants had higher chlorophyll *a* and *b* content at low intensity (150 µmol photons m^-2^ s^-1^) of white light. Under K deficiency, at 450 µmol photons m^-2^ s^-1^, chlorophyll *a* and chlorophyll *b* increased significantly in the female plants compared to that in the male plants. (Fig. 2 A, B and SM Table S4).

**Figure 2.**
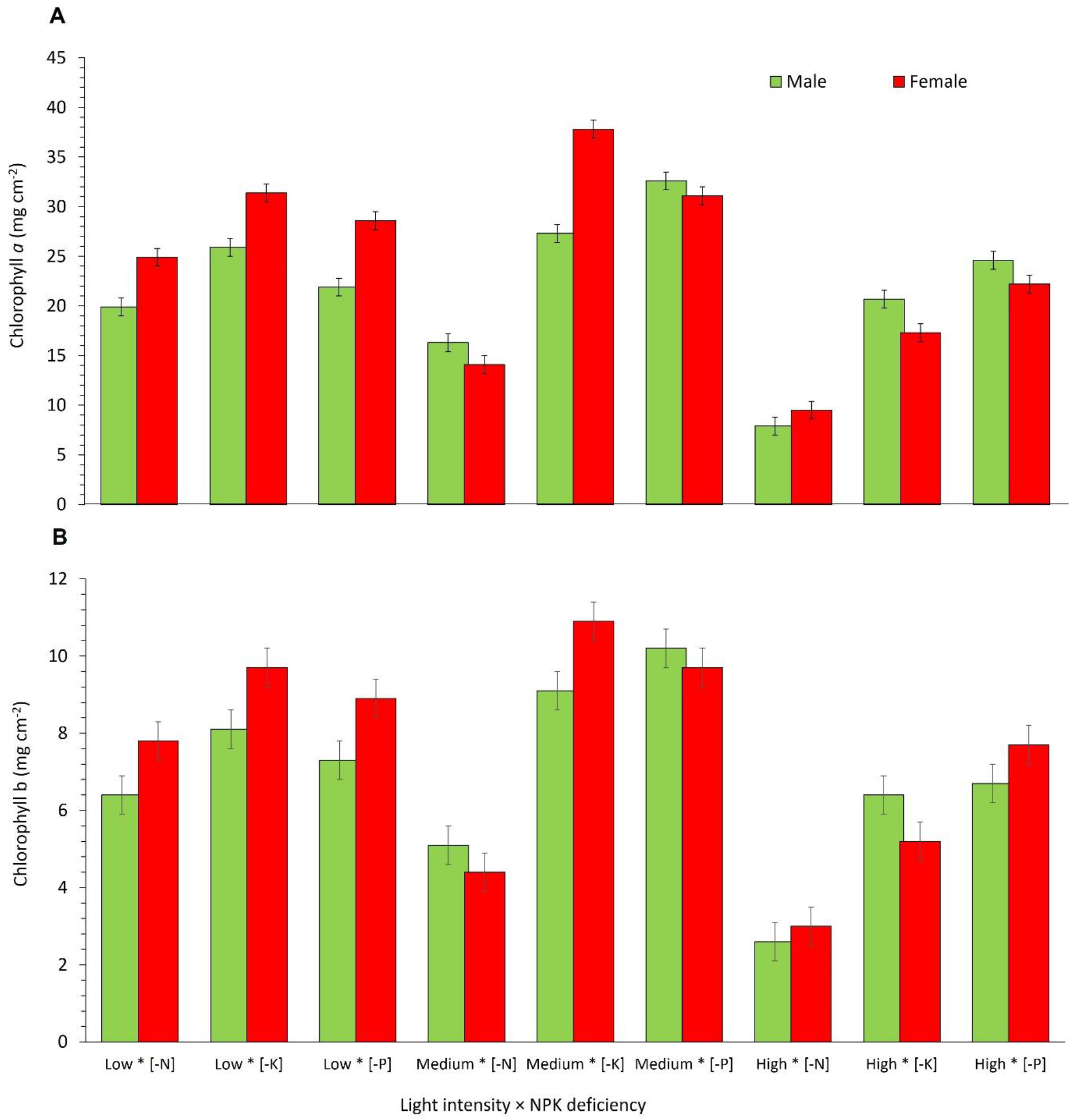
(A) Leaf chlorophyll *a* content, and (B) leaf chlorophyll *b* content of male and female *Amaranthus palmeri* plants as affected by white light intensity × NPK deficiency averaged across 5 sampling times throughout the experimental period; Values of chlorophyll *a* and chlorophyll *b* content for male and female *A. palmeri* plants on the Y-axis stand for the interaction of white light intensity × NPK deficiency shown on the X-axis. Low, Medium, and High light intensities refer to 150, 450 and 1300 µmol photons m^-2^ s^-1^, respectively; [-N], [-K], and [-P] refer to NPK deficiency. Vertical bars represent Least Significant Difference at *a*=0.05.

However, no difference in the chlorophyll content was recorded between the female and the male plants, grown under P deficiency at medium (450 µmol photons m^-2^ s^-1^) white light intensity. The content of chlorophyll *a* and *b* in female compared to male plants, under PK deficiency and high white light intensity (1300 µmol photons m^-2^ s^-1^) showed a significant decrease (P = 0.007 for chlorophyll *a*, and P= 0.006 for chlorophyll *b*; see Fig. 2 A, B and SM Table S4).

### Effects of white light intensity on chlorophyll α and chlorophyll *b* in *Amaranthus palmeri* male and female plants

Female plants, grown at 150 µmol photons m^-2^ s^-1^ of white light, had higher chlorophyll α and chlorophyll *b* content at their earlier life stages (i.e., 14, 21 and 28 days after treatment initiation- DAT; see Fig. 3 A, B and SM Table S4). However, with increasing time (i.e., 35 and 42 DAT), chlorophyll α and chlorophyll *b* content, averaged across NPK deficiencies, was higher (P<0.0001) in male compared to female plants (Fig. 3 A, B), grown at 1300 µmol photons m^-2^ s^-1^ of white light. Gradual increases in white light intensity led to decreases in chlorophyll *a* and *b* content, especially at high light intensity, in both male and female *A. palmeri* plants.

**Figure 3.**
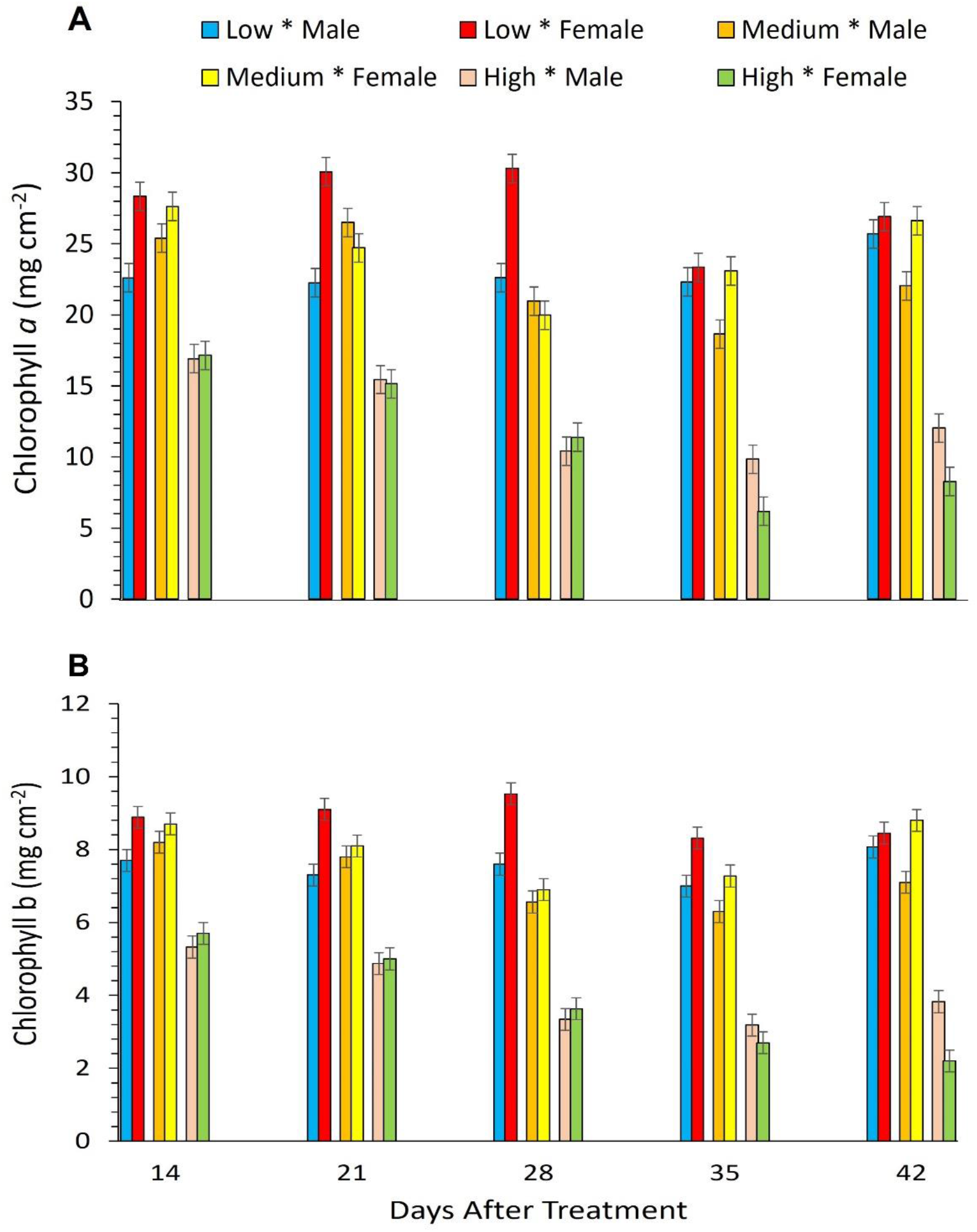
(A) Leaf chlorophyll α content, and **(B)** leaf chlorophyll *b* content in male and female *A. palmeri* plants at 3 different white light intensities as measured throughout the experimental period (DAT) and averaged across NPK deficiency treatments. Low, Medium, and High refer to 150, 450 and 1300 µmol photons m^-2^ s^-1^ (of light) respectively for male or female plants; vertical bars represent Least Significant Difference at *a*=0.05.

Consequently, the chlorophyll *a*/*b* (Chl α/*b*) ratio was reduced (P=0.0002) 28 DAT onwards in both male and female *A. palmeri* plants, compared to plants that were kept at lower intensities of white light (150 and 450 µmol photons m^-2^ s^-1^) (Fig. 4 and SM Table S4). Furthermore, the Chl α*/b* ratio at 38 and 45 DAT at high light intensity (1300 µmol photons m^-2^ s^-1^) was significantly greater (P=0.0002) in the male compared to that in the female plants (Fig. 4 A, B and SM Table S4).

**Figure 4.**
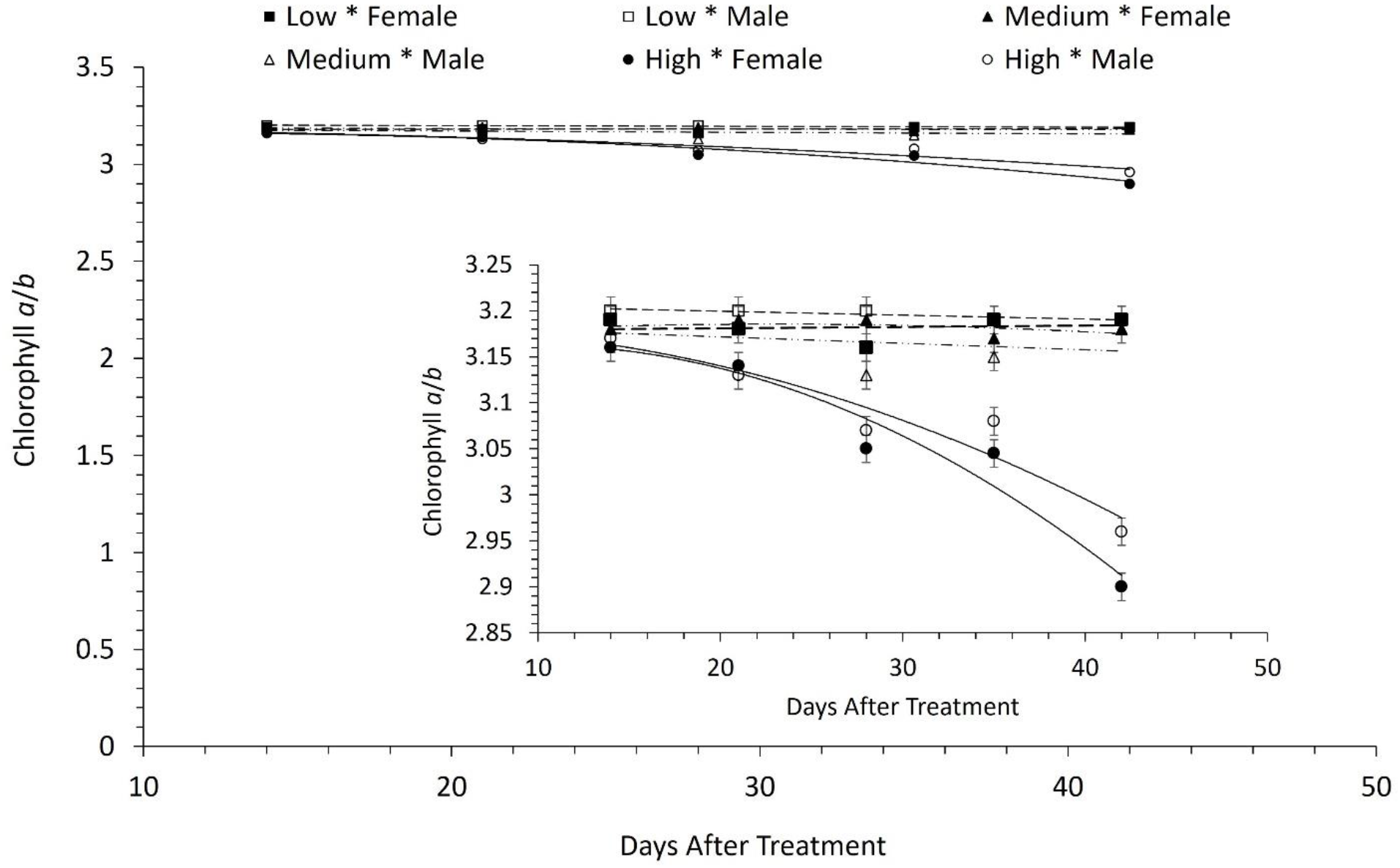
Effects of white light intensity on leaf chlorophyll *a*/b ratio as measured throughout the experimental period (DAT) for *Amaranthus palmeri* male and female plants and averaged across NPK deficiency treatments. The Y-axis, in the inset figure, has been rescaled to improve the readability of the figure. Squares, in the inset figure, represent the chlorophyll *a*/*b* (Chl *a*/*b*) ratio in male plants (open squares), and in female plants (closed squares) both under low light intensity (150 μmol photons m^-2^ s^-1^) (see dashed line). Open triangles are for the data from the male plants, and closed triangles are for female plants, both under medium light intensity (450 μmol photons m^-2^ s^-1^) (long dashed line with dots). The Chl *a*/*b* ratio under high light intensity (1300 μmol photons m^-2^ s^-1^) (solid line) is shown for both the male (male sign) and the female (female sign) plants. Vertical bars represent LSD=Least Significant Differences at *a*=0.05.

The chlorophyll content in untreated *A. palmeri* plants throughout the experimental period showed no significant differences between the male and female plants. Chlorophyll *a* content was 31.7 and 32.5 mg cm^-2^ (P=0.722) and chlorophyll *b* was 9.97 and 10.25 mg cm^-2^ (P=0.712) for male and female plants, respectively. In addition, no interaction between the untreated male and female plants was observed (P=0.406 and P=0.478 for chlorophyll *a* and chlorophyll *b*, respectively) when the sampling time was treated as a fixed variable. Likewise, no significant difference between the untreated male and female *A. palmeri* plants was observed for Chl *a*/*b* ratio (P=0.351) when the sampling time was treated as a fixed variable.

### High white light intensity reduces the operating capacity of PS II in the female plants of *Amaranthus palmeri*

Analysis of chlorophyll fluorescence parameters (F’_S_, F’_M_) and Φ_PSII_ revealed an interaction between the white light intensity, the sex of *A. palmeri* and the sampling time (P<0.0001) (SM Table S5). We observed significant differences between the male and the female plants at high intensity (1300 µmol photons m^-2^ s^-1^) of white light, 28 DAT onwards in both steady state chlorophyll fluorescence (F’_S_) (P=0.09) and maximum fluorescence (F’_M_) (P=0.03) (Fig. 5 A, B and SM Table S5).

**Figure 5.**
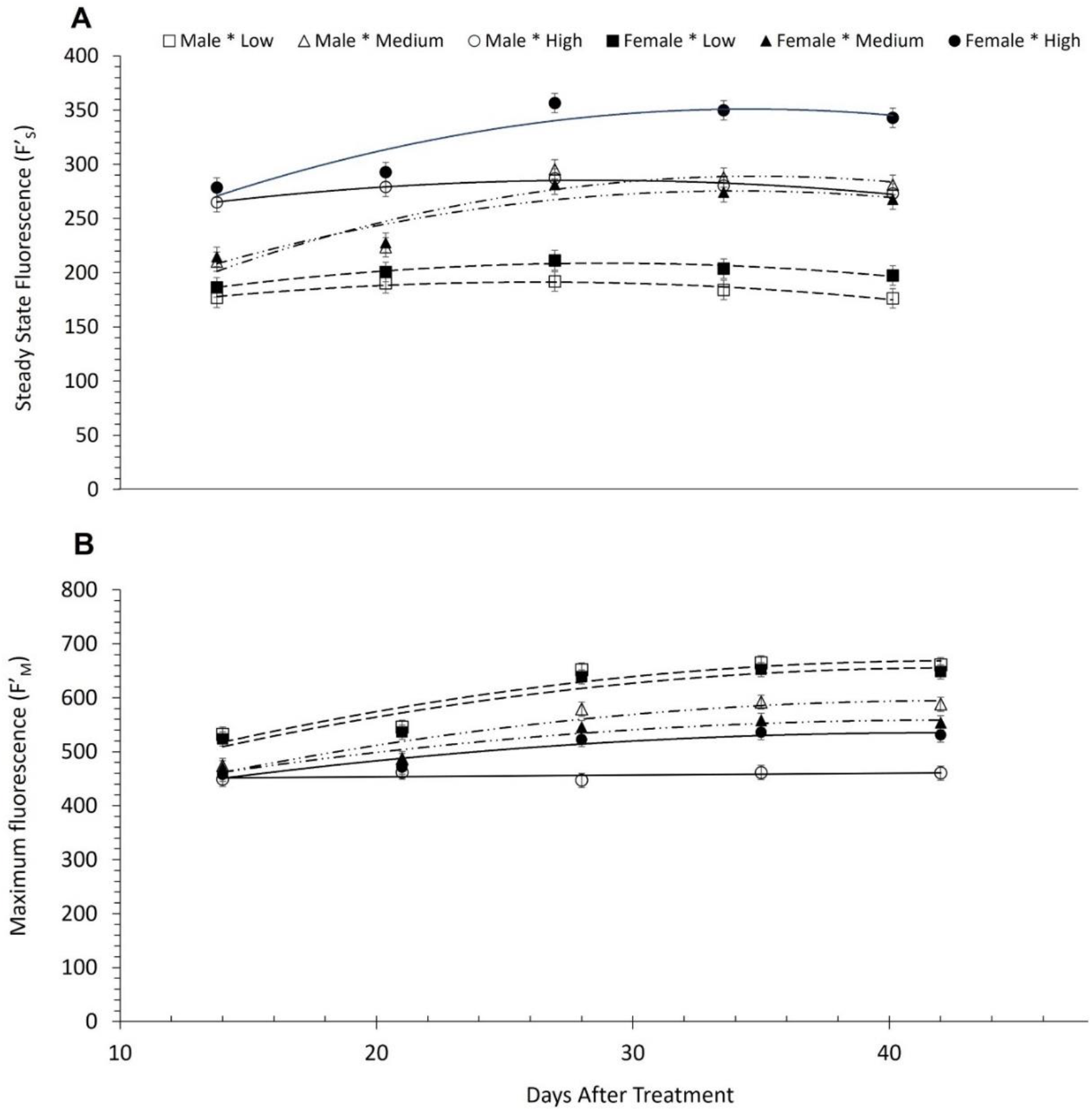
**(A)** Steady state chlorophyll *a* fluorescence (F’_S_) for leaves; and (**B**) Maximum chlorophyll *a* fluorescence (F’_M_) from the leaves of male and female *A. palmeri* plants, grown at 3 different white light intensities, measured throughout the experimental period (DAT) and averaged across NPK deficiency treatments. Open and closed squares: Male and female F’_S_ and F’_M_ respectively under low light intensity (150 μmol photons m^-2^ s^-1^) (dashed line); open and closed triangles: Male and female F’_S_ and F’_M_ respectively under medium light intensity (450 μmol photons m^-2^ s^-1^) (long dashed line with dots); open and closed circles: Male and female F’_S_ and F’_M_ respectively under high white light intensity (1300 μmol photons m^-2^ s^-1^) (solid line). Low, Medium, and High refer to 150, 450 and 1300 µmol photons m^-2^ s^-1^ respectively, used for growing male or female plants; vertical bars represent Least Significant Difference at *a*=0.05.

Furthermore, male plants had a greater (22.5%) Φ_PSII_ value (P=0.04) at 1300 µmol photons m^-2^ s^-1^, 35 and 42 DAT compared to female plants. However, both male and female plants showed lower Φ_PSII_ values at higher light intensity (1300 µmol photons m^-2^ s^-1^) in comparison to those that were kept at 150 and 450 µmol photons m^-2^ s^-1^ of white light. More specifically, Φ_PSII_ values were 44.5 and 66.1% lower for male and female plants respectively at high light intensity compared to Φ_PSII_ values at low white light intensity. Similarly, the Φ_PSII_ values at 1300 µmol photons m^-2^ s^-1^ were 26 and 55.3% lower for male and female plants respectively at high light intensity compared to medium light intensity) (Fig. 6 and SM Table S5).

**Figure 6.**
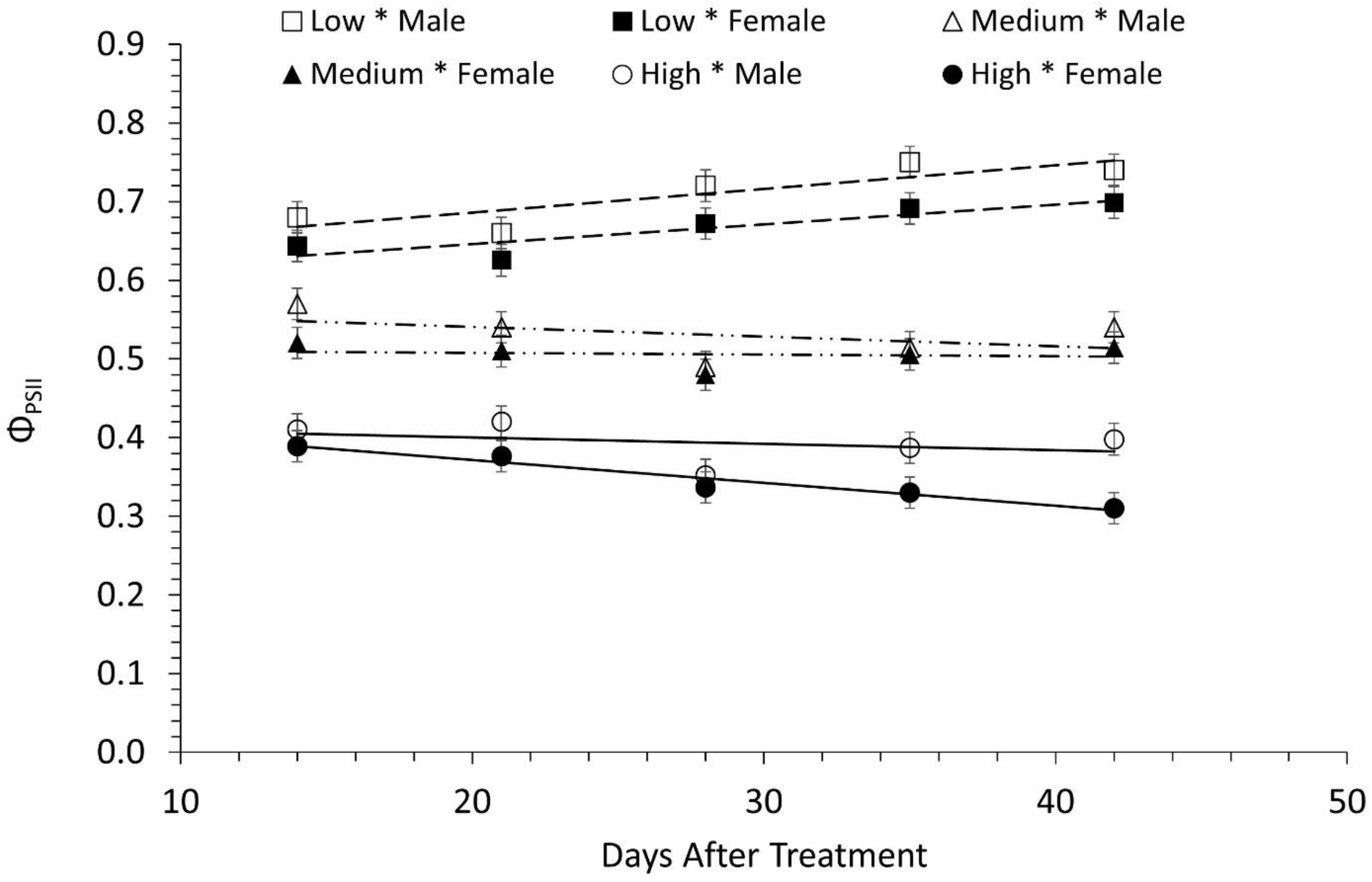
Photochemical operation efficiency of PSII (Φ_PSII_) in male and female *A. palmeri* plants, grown at 3 different white light intensities measured throughout the experimental period (DAT) and averaged across NPK deficiency treatments. Open and closed squares: Φ_PSII_ of male and female plants, grown under low light intensity (150 μmol photons m^-2^ s^-1^) (dashed line); open and closed triangles: Φ_PSII_ (of male and female plants) under medium light intensity (450 μmol photons m^-2^ s^-1^) (long dashed line with dots); open and closed circles: F’_S_ and F’_M_ (of male and female plants) grown under high white light intensity (1300 μmol photons m^-2^ s^-1^) (solid line). Vertical bars represent Least Significant Difference at *a*=0.05.

Analysis of untreated controls of *A. palmeri* showed no differences between male and female plants for chlorophyll fluorescence parameter F’_S_ (P=0.613) or between *A. palmeri* sex × sampling time (P=0.056). Likewise, no differences were obtained for F’_M_ between the male and the female plants (P=0.992) or when these parameters were analyzed against the sampling time (P=0.586). F’_S_ values for female and male *A. palmeri* plants, averaged across sampling times, were 189.8 and 193.4 (P=0.613) whereas F’_M_ measured at 495.1 and 495 (P=0.992). Likewise, no differences were obtained for Φ_PSII_ between the male and female plants (P=0.491), or when the values of Φ_PSII_ of both male and female plants were analyzed against the sampling time (P= 0.109). Φ_PSII_ of untreated *A. palmeri* male and female plants was 0.72 and 0.71 (P=0.491) respectively.

### Effects of NPK deficiency on F’_M_ and Φ_PSII_ parameters

Significant differences between male and female plants were observed in F’_M_, and Φ_PSII_ values when they were under mineral deficient conditions (SM Table S5). F’_M_ increased at 14 DAT onwards in both male and female plants independent of mineral deficiency (Table 2). However, the female plants had significantly higher F’_M_ values, 5.5% higher F’_M_, under N deficiency compared to those in the male plants 42 DAT. In addition, the F’_M_ values for the male plants, when they were grown under N deficiency, were lower compared to the corresponding values under P and K deficiency, particularly 28 DAT onwards, but interestingly, this was not the case for the female plants (Table 2). Further, higher Φ_PSII_ values, ranging between 7.3% and 10.5%, were obtained for the male compared to female plants 28 DAT onwards under P deficiency, and to lesser extent under K and N deficiency (Table 2).

**Table 2.**
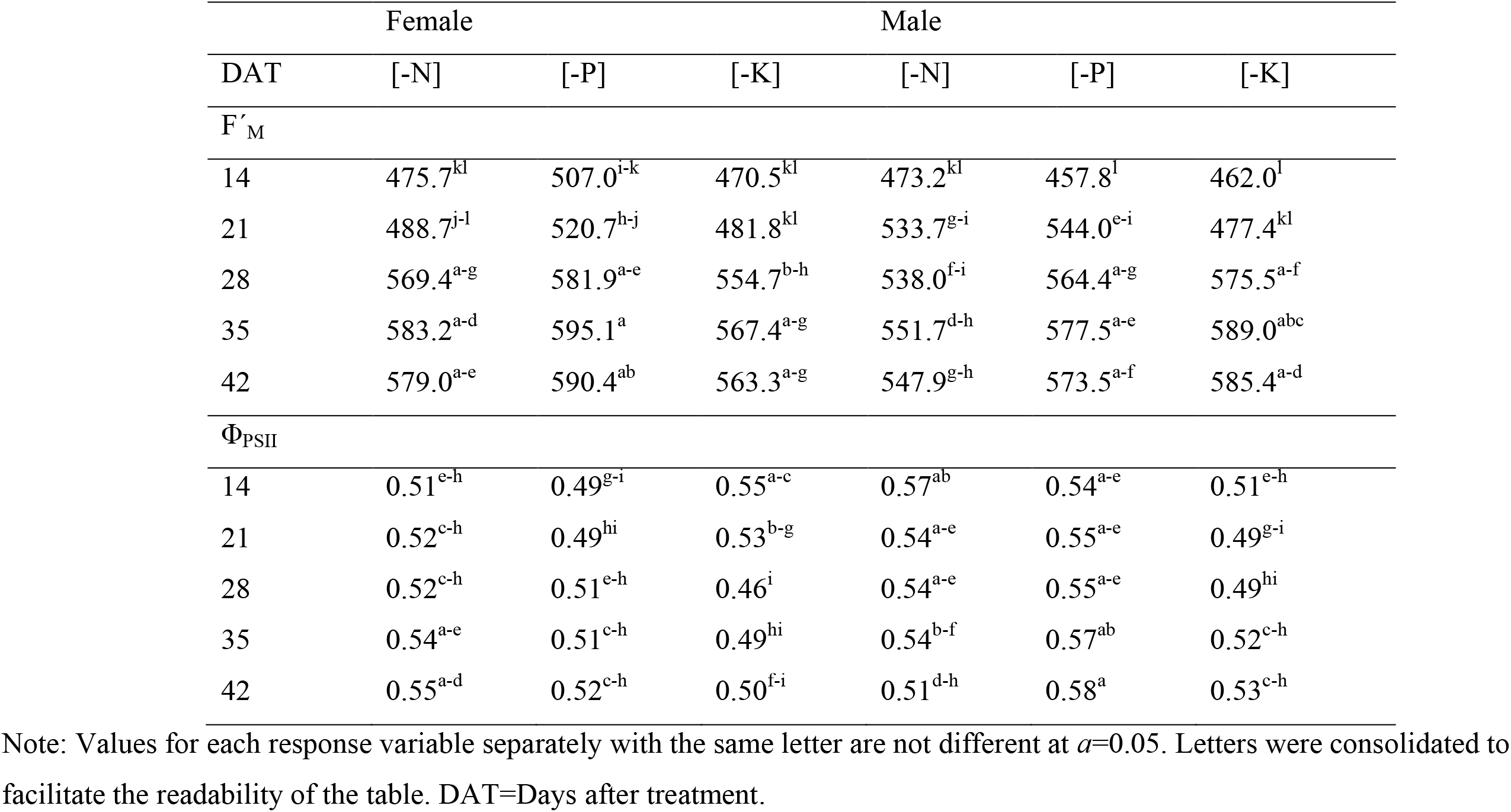
Effects of NPK deficiency on *A. palmeri* sex on values of F’_M_ and Φ_PSII_ throughout the experimental period

## Discussion

Secondary sexual dimorphism is the differentiation of traits between the male and the female plants, which is not directly linked to gamete production (Dudley, 2006). In this research, we examined the secondary sexual dimorphism related to functional characteristics, such as the mineral content, the dynamics of changes in chlorophyll content, and chlorophyll fluorescence parameters, related to photosynthesis, in both male and female plants of *A. palmeri*.

### *I*nter- and intrasexual differences in mineral content due to abiotic stress

Changes in the content of minerals in plant organs are related to dry matter accumulation (Hocking and Pate, 1976). Female plants accumulate higher amount of minerals in the stems (including inflorescences) than in the leaves in contrast to the male plants which accumulate relatively equal amounts in both leaves and stems (Table 1). The accumulation of mineral elements, especially N, P, K, and Mg in the stem and inflorescences, is particularly important for seed filling (Hocking and Pate, 1976; Ding et al., 2005). Further, the extent of mineral accumulation in plant organs depends upon the interrelationship between the “sink-size” e.g., the biomass produced or the number of seeds with the “source-size” e.g., leaf area or specific leaf area (Marschner, 1983), but also whether the plants and seeds are inadequately supplied with mineral nutrients (Loneragan et al., 1976), as shown in this paper. The NPK deficiency was found to be the most important factor affecting the mineral content in the stems (including inflorescences) and in the leaves of male and female *A. palmeri*; this is especially the case for the stems (incl. inflorescences) of female plants (Table 1 and SM Table S1). Therefore, it is reasonable to suggest that the higher accumulation of minerals in the stems (incl. inflorescences) compared to the leaves in the female pants are related to the reproductive demand for resources, a sex-specific characteristic in dioecious species, as already noted by Nowak-Dyjeta et al. (2017). The female plants are more efficient in gathering or using certain resources when compared to the male plants (Lei at al., 2017). Natural selection in dioecious species, that initiated the evolution of sexual dimorphism (Brock et al., 2017), allows *A. palmeri* female plants to reproduce successfully under resource limiting environments by exploiting the resources more efficiently than those in the male plants. No effect of light intensity on stem or leaf mineral content of *A. palmeri* sex was observed in our research (SM Table S1). Currently, a controversy exists about the effects of light intensity on the nutrient content of plant organs. Li et al. (Li et al., 2020), for example, have shown that low white light supply significantly decreases the content of leaf N in soybean [*Glycine max* (L.) Merr] plants, whereas Bouma (1983) has shown that when plants were grown at low white light intensities, P content was higher in all parts of the subterranean clover (*Trifolium subterraneum* L.) plant. Thus, the mineral content allocation and accumulation in plant organs, exposed to a range of light intensity, is a species-specific storage strategy to cope with light stress. Korres et al. (2017) have shown that female plants of *A. palmeri* respond to shading by stem elongation, whereas the male plants respond by increasing specific leaf area.

In this current paper, we have clearly observed sex-specific intra- and intersexual correlations between leaf and stem mineral content (Fig. 1 and SM Tables S2, S3). The synergistic interaction between N and K, two important minerals in regulating leaf photosynthesis, has been well documented in other plants (Hou et al., 2019); however, such association is absent in *A. palmeri* female plants (Fig. 1 A, C, SM Table S2). In contrast, Ca and Mg, two important elements for determining the structural rigidity of the cell wall (Anonymous, 2005) and chlorophyll production (Hao and Papadopoulos, 2004), respectively, have been shown here to be positively correlated in leaves of both *A. palmeri* sexes, but only in the stems of the male plants (Fig. 1, D, SM Tables S2, S3). The positive correlation between Ca and Mg indicates a synergistic interaction between these minerals in both male and female plants. Lopez-Lefebre et al. (2001) showed that Ca exerts a positive effect on N assimilation by activating enzymes responsible for N assimilation. However, no such relationship was observed in our current research (Fig. 1, SM Tables S2, S3) suggesting a species specificity of Ca-N association. A negative correlation was observed between K and Ca in the leaves of the male, but not in the female plants (Fig. 1 A, B, SM Tables S2, S3) indicating an antagonistic interaction between these two minerals in male plants. Iron, an essential mineral for photosynthesis and chlorophyll synthesis (Wu et al., 2019), is another mineral that is shown here to be associated negatively with K in both leaves and stems of male plants (Fig. 1 B, D, SM Tables S2, S3). Wu et al. (2019) reported that potassium affects iron translocation, whereas Bolle-Jones (1955) suggested that iron may impede the translocation of potassium.

### Effects of NPK deficiency and white light intensity on chlorophyll content in male and female *Amaranthus palmeri* plants

Nitrogen is an essential component of chlorophyll (Ding et al., 2005), the content of which decreased in both male and female plants under nitrogen deficiency and when these plants were grown at high light intensity. Our results affirm the findings of Ding et al. (2005) and Li et al. (2020) demonstrating decreases in chlorophyll *a* and *b* content under N deficiency. Research on the effects of abiotic stress on sexually related secondary traits, including chlorophyll content, has been, in the past, conducted mostly as mono-factorial experimental designs. However, plants are exposed to multidimensional environmental changes that occur concurrently, as in our paper. Therefore, it is rational to suggest that increased light intensity which can potentially damage or even destroy chlorophyll *a* and *b* (Glime, 2017; Feng et al., 2019) in conjunction with N deficiency, have possibly caused decreases in the content of both chlorophyll *a* and *b* in both sexes. On the contrary the effects of P and K deficiencies have been shown to be less deleterious on the content of chlorophyll *a* and *b* compared to that by N deficiency; chlorophyll *a*, for example, shows a fluctuating response between male and female plants under P and K deficiencies and different white light intensities, indicating the importance of N deficiency on chlorophyll content. However, at high light intensity (i.e., 1300 µmol photons m^-2^ s^-1^) and PK deficiencies, the male plants had greater chlorophyll *a* content compared to the female plants (Fig. 2 A). A similar effect was observed for chlorophyll *b* under high light intensity (i.e., 1300 µmol photons m^-2^ s^-1^) and K deficiency (Fig. 2 B). On the contrary, female plants exhibited greater chlorophyll *a* and *b* content at low and medium light intensities (150 and 450 µmol photons m^-2^ s^-1^) and N, P or K deficiency (Fig. 2 A, B).

Feng et al. (2019) found that light intensities ranging between 100-500 µmol photons m^−2^ s^−1^ resulted in gradual increases in both chlorophyll *a* and *b* content in soybeans. However, the light intensity used by Feng et al. (2019) was much lower compared to the light intensity of 1300 µmol photons m^−2^ s^−1^ used in our research here. On the other hand, Marshall and Proctor (2004) found that light intensities of 1000 µmol photons m^−2^ s^−1^ led to low chlorophyll content but high Chl *a*/b ratio in bryophytes, whereas, in barley leaves, Zivcak et al. (2014) observed the opposite between low and high irradiance ranging from 100 to 1200 µmol photons m^−2^ s^−1^. Nevertheless, decreases in the amount of chlorophyll, observed under conditions of stress such as high light intensity (Glime, 2017), have been attributed either to disruption in chlorophyll synthesis, increased degradation of chlorophyll due to higher activity of the enzyme chlorophyllase or due to oxidative damage with consequent destruction of chloroplasts (Hortensteiner, 1999; Zhang et al., 2016). In our work, higher content of chlorophyll *a* and *b* was observed in the male plants, at high light intensity (i.e., 1300 µmol photons m^-2^ s^-1^) at 35 and 42 DAT compared to that in the female plants, which showed greater chlorophyll *a* and *b* content at low light intensities up to 28 DAT (Fig. 3 A, B). These effects can be attributed to ontogenetic differences between *A. palmeri* sexes. Korres et al. (2017), for example, have reported a greater leaf area and specific leaf area values in the male than in the female plants at high light (i.e., 1300 µmol photons m^-2^ s^-1^) intensity, and under NPK mineral deficiency. Therefore, it is reasonable to suggest that increases of chlorophyll content in the male plants at high light intensity (i.e., 1300 µmol photons m^-2^ s^-1^) towards maturity reflects intersexual differences in secondary sex-specific ontogenetic traits such as leaf area and specific leaf area. Chl *a* affects the photosynthetic activity in plants (Sestak, 1996; Papageorgiou and Govindjee, 2004); thus, the tendency towards a higher content of chlorophyll *a* and *b* in the male plants at high light intensity (Fig. 3 A, B) might partially explain their increased photosynthetic capacity mentioned below.

Grieco et al. (2012) have used the Chl *a*/*b* ratio as a parameter to monitor the acclimation processes of *Arabidopsis thaliana.* According to Anderson and Andersson (1998), differences in this ratio reflect modifications in the stoichiometry of the photosynthetic pigment protein complexes. Therefore, lower Chl *a*/*b* ratio in female plants at 42 DAT (Fig. 4) might reflect modifications of stoichiometry between Chl *a* and Chl *b* as an adaptation mechanism, to compensate for decreases in their photosynthetic capacity (Friedland, 2019) under high light intensities (Glime, 2017). Further, Spundova et al. (2003) explained decreases in Chl *a*/*b* ratio as counteractive mechanism for photooxidative damage and consequent losses in energy utilization efficiency (Dinc, 2012). Decreases in energy utilization efficiency are known to result in the accumulation of excess excitation energy in the photosynthetic chain (Ramel et al., 2012) and ultimately in the over-production of reactive oxygen species (ROS) (Foyer, 2018).

### Effects of white light intensity and NPK deficiency on the operating efficiency of PSII in male and female *Amaranthus palmeri* plants

Chlorophyll fluorescence is a reliable tool to study and quantify photosynthesis and responses of plants to environmental changes and abiotic stresses such as mineral deficiencies or light restricted environments (Murchie and Lawson, 2013; Kalaqji et al., 2016; Lichtenthaler et al., 2005).

Steady state fluorescence (F’_S_), used in our research, is the fluorescence emission from a light- adapted leaf induced by a non-saturating irradiation (Lichtenthaler et al., 2005). Female plants of *A. palmeri* show significantly higher F’_S_ values under high light intensity, especially 28 DAT onwards, compared to the male plants, indicating a lower photosynthetic performance of the former, given the competitive nature between the two processes i.e., chlorophyll fluorescence and photosynthetic efficiency (Govindjee, 2004). On the other hand, F’_M_ describes the state of PSII when all its reaction centers are closed or reduced, i.e., the electron transport carrier Q_A_ (the first bound plastoquinone) is fully reduced and cannot pass electrons to other components, that are also fully reduced (Papageorgiou and Govindjee, 2004). The higher values of F’_M_ in female *A. palmeri* plants, under high light intensity, as time progresses (i.e., 28 DAT onwards) (Fig. 5 A, B) are possibly caused by decreases in the content of Chl *a*, and Chl *b,* and in the Chl *a*/*b* ratio, mentioned above, especially at the later stages of the life cycle of the plant. Chlorophyll *a* and *b* content reduction is related to decreased PSII capacity due to decreased electron transport (Murchie and Lawson, 2013), hence energy transfer. Changes in energy transfer coherence, possibly due to changes in the orientation of the transition dipoles of Chl molecules or the structure of PSII apparatus (Kalaji et al., 2016), due to abiotic stress in both sexes of *A. palmeri*, might have spurred decreases in Φ_PSII_ (Fig. 6). Φ_PSII_, the quantum yield of PSII in the light- adapted state, is related to the operational efficiency of PSII (Genty et al., 1992), hence to the photosynthetic efficiency under different light intensities (Baker, 2008). Several studies have shown a direct or indirect relationship between Φ_PSII_ and chlorophyll content in leaves (Glime 2017; Lichtenthaler et al., 2005; Baker, 2008; Kumagai et al., 2009). In support of the above hypothesis, Φ_PSII_ was found to be positively correlated with chlorophyll content in both male and female *A. palmeri* plants (R^2^=0.313 and 0.272 for male and female plants respectively) (Fig. 7). Further, a similar trend was observed between Φ_PSII_ and chlorophyll *a* and *b* in both *A. palmeri* male and female plants.

**Figure 7.**
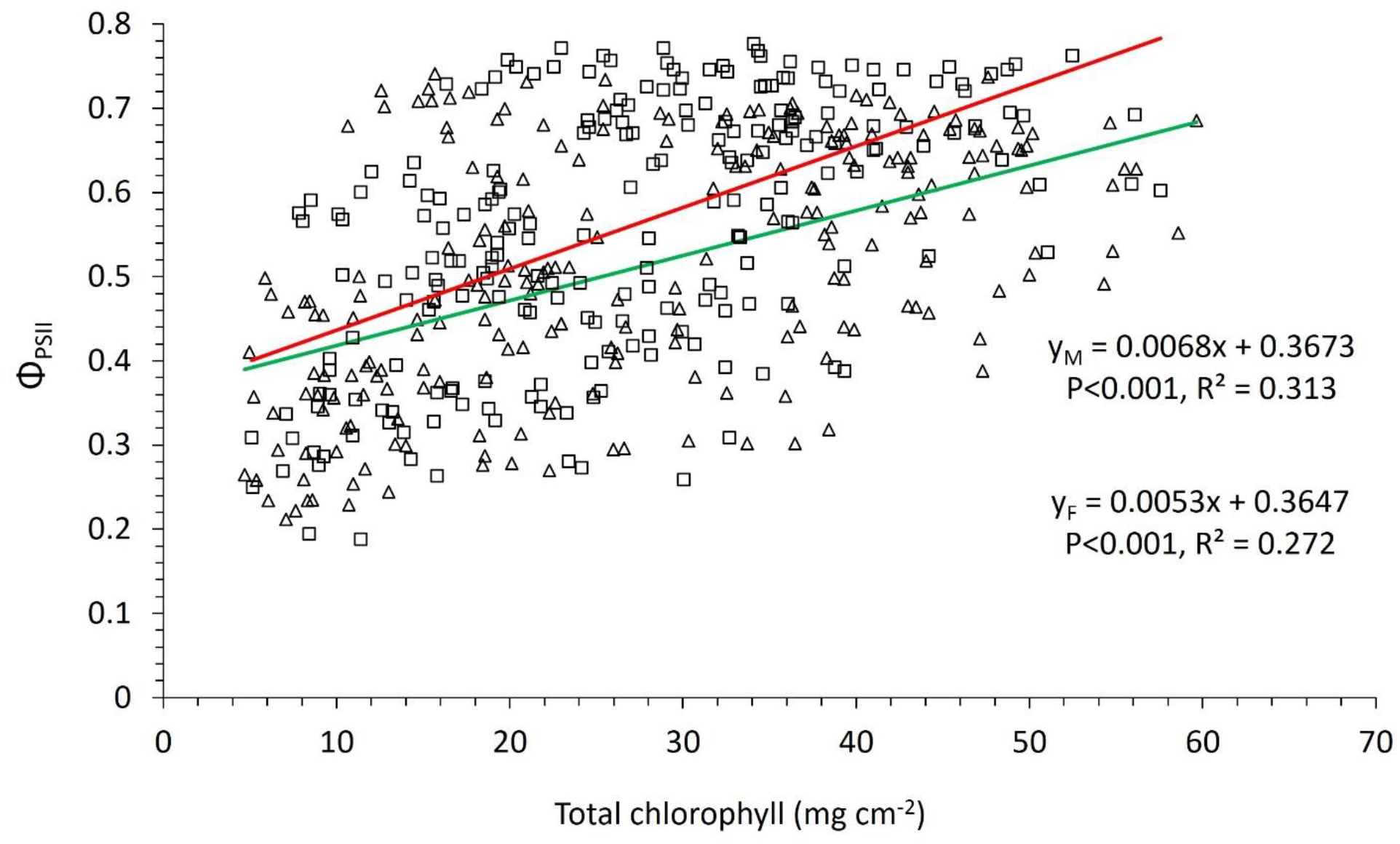
Correlation between the operating efficiency of PSII (Φ_PSII_) and the total leaf chlorophyll content in male (open squares) and female (open triangles) *A. palmeri* plants. Y_M_ and Y_F_ in the equations denote Φ_PSII_ for male and female *A. palmeri* plants, respectively, and *x* denotes the total leaf chlorophyll content. A red colored curve is fitted for data from male plants, and a green colored curve is for data from female plants.

The lower Φ_PSII_ values, especially at the mature life stages of the female plants under high white light intensity (Fig. 6) seemed to be caused by a low PSII operating capacity, as suggested by the low Chl *a*/*b* ratio (Bertamini and Nedunchezhian, 2003) and observed at high white-light intensity in this research. Surprisingly, the electron transport rate (ETR) and total chlorophyll content are negatively correlated (Fig. 8), especially in female plants, with R^2^=0.362 vs R^2^=0.099 for female and male plants respectively (same trends were observed between ETR and chlorophyll *a* and chlorophyll *b*). This is an indication of a negative energy feedback that denotes a photoinhibition symptom, despite the positive correlation between Φ_PSII_ and chlorophyll content, which causes reduction in the operating efficiency of *A. palmeri* PSII (Roach et al., 2014), especially in the female plants. Further research is needed to find whether photoinhibition results from photodamage or it is a regulatory and protective adjustment to abiotic stress related to mineral deficiency and white light environment. It is also imperative to quantify these effects in relation to the mechanisms of non-photochemical quenching, including thermal dissipation and photosynthesis state transitions. Finally, investigating the dynamics between chlorophyll content and PSII operation efficiency will expand our knowledge in up- and down-stream regulatory processes in photosynthesis.

**Figure 8.**
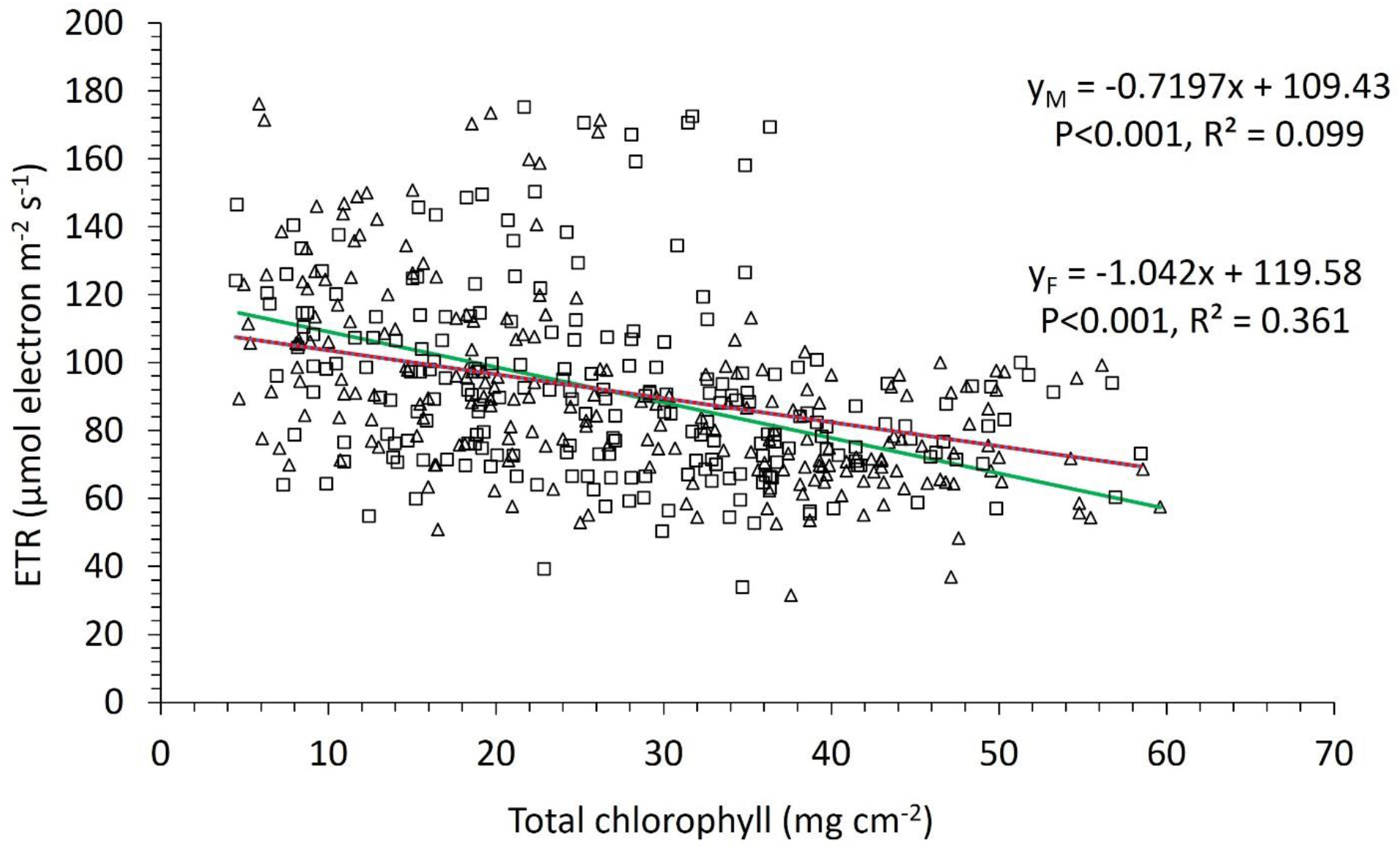
Correlation between electron transport rate (ETR) and total leaf chlorophyll content in male (open squares) and female (open triangles) *A. palmeri* plants. Y_M_ and Y_F_ in the equations denote ETR for male and female *A. palmeri* plants respectively, and *x* stands for the total leaf chlorophyll content. A red colored curve was fitted for data (from male plants) and a green colored curve was fitted for data from female plants.

The lower Φ_PSII_ values of the female *A. palmeri* plants, compared to that of the male plants under P and to lesser extent under K and N deficiencies (Table 2) indicate the lower PS II operating efficiency of the former. Korres et al. (2017) reported a lower leaf area and specific leaf area in female compared to male *A. palmeri* plants, under NPK deficiency and high light intensity treatment, which was accompanied by the presence of albino leaves (SM Fig. S2,), an indication of lower chlorophyll content. Further, our results on *A. palmeri* plants agree with the results on nutrient deficient *Populus* plants, where the photosynthetic efficiency of male plants was higher than that of the female plants (Zhang et al., 2004). Thus, our results on the differences in male and female *A. palmeri* plants are of general importance to overall plant biology. Furthermore, our research has revealed new information on the ontogenetic and physiological secondary traits that make *A. palmeri* one of the most successful competitors in nature.

We suggest that the dioecious nature of the species has weaknesses based on which we can adjust our weed management approaches by manipulating the microclimate, e.g., light/dark conditions or mineral availability at a field scale, without relying solely on the herbicides for its control. In addition, the exogenous or endogenous stressful photodynamic treatments with the aim of decreasing the amount of chlorophyll through natural photosensitizers, for example porphyrins, merits further investigation. Rebeiz et al. (1984) reported that dicotyledonous weeds including *Amaranthus retroflexus*, another species of the Amaranthaceae family, are susceptible to photodynamic treatments, especially when used in monocotyledonous crops such as maize (*Zea mays* L.), where *A. palmeri* is a major problem.

## Materials and Methods

### Plant material, experimental design, and treatment arrangements

*Amaranthus palmeri* seeds were collected from a field at the University of Arkansas, Fayetteville, AR. Cuttings from 50 different F1-generation plants of each sex, as described by Korres *et al*. (2017), were placed under NPK deficient conditions and different white light intensities. Each NPK deficiency treatment contained only 10% N, P, or K of the standard N (91.4 g NH_4_NO_3_ L^−1^), P (40.3 g NaH_2_PO_4_ ×2H_2_O L^-1^) or K (71.4 g K_2_SO^4^ L^−1^) stock solutions, (cf. Ref. Yoshida et al., 1976). For low, medium and high light we used 150, 450 and 1300 μmol photons m^−2^ s^−1^ of white light respectively. The experimental design was based on a randomized complete block arrangement with white light intensity (one growth chamber for each white light intensity) as blocking treatment and N, or P, or K deficiency (henceforth NPK deficiency) for both male and female *A. palmeri* cuttings as randomized treatments within each block. More specifically, for each white light-intensity, each of the 20 *A. palmeri cuttings* (10 cuttings from the male parents and 10 cuttings from the female parents) were exposed to NPK deficient environment. Also, 10 plants from each *A. palmeri* sex, receiving full nutrition, were left to grow under greenhouse conditions, and used as controls. The above set of treatments was conducted twice. A detailed description of plant material preparation, experimental design and treatments is provided in supplemental material (SM, Materials and Methods).

### Determination of mineral content in the stems and leaves of *A. palmeri* male and female plants

At final harvest time (i.e., before the dormant stage), leaves and stems (the latter including inflorescence) taken from five randomly selected plants were dried in a forced-draft oven at 70 °C for 48 h and mechanically ground prior to the determination of their mineral content, which was conducted by the Agriculture Diagnostic Laboratory, University of Arkansas, Fayetteville, USA. The determination of plant leaf and stem mineral concentrations was made after digestion with a mixture of HNO_3_/peroxide digest using hot block “digest” and inductively coupled Agron plasma (ICP) emission spectrometry, as described by Jones and Case (1990). Percent N was determined by combustion as described by Campbell (1992). To facilitate statistical analysis, all mineral concentrations were expressed in relation to the leaf (and stem) dry weight as micrograms per g of leaf dry weight.

### Chlorophyll content

Chlorophyll *a* and *b* of both male and female *A. palmeri* plants was assessed by taking five measurements from five fully expanded, light-exposed leaves in the upper half of the main stem (the same leaves were used for chlorophyll fluorescence measurements; see below) from five randomly selected plants per treatment, using a handheld SPAD-502 (Minolta Camera, Osaka, Japan). The estimation of total chlorophyll, chlorophyll *a* and *b* in mg cm^-2^ was based on the calibration equations as developed by Richardson *et al*. (2002) and Porra *et al*. (1989).

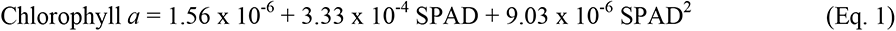

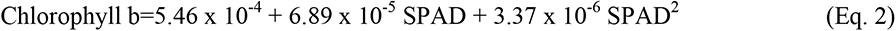

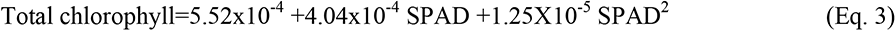

where SPAD=SPAD units.

### Chlorophyll fluorescence

The photochemical efficiency (Φ_PSII_) of Photosystem II (PSII) complex in the light-adapted state was estimated via chlorophyll fluorescence, as described by Baker (2008); here we used OS5p modulated fluorometer (Opti-Science, Tyngsboro, MA) for these measurements. For each treatment, five light-adapted, fully expanded young leaves, from the upper half of the main stem of 5 randomly selected *A. palmeri* plants, were chosen. These measurements were made weekly for both the experimental runs from the leaves of the same randomly chosen plants at the beginning of the experiment. Data was recorded 5 h after the initiation of the daily 14 h white light intensity treatment, i.e., in the photoperiod cycle, when the leaves were in a fully light- adapted state. Using exciting light of 1800 µmol photons m^-2^ s^-1^, we measured changes in chlorophyll fluorescence: F*’q* or ΔF=F’_M_-F’_S_, where F′_S_ is the steady state fluorescence in the light-adapted state, and F′_M_ is the maximum fluorescence induced by a saturating light pulse immediately after F′_S_; note that F*’q* is a measure of the photochemical quenching of chlorophyll fluorescence by open PSII centers (Baker, 2008). In light-adapted leaves at high actinic light history, with all PSII reaction centers closed measurement of F′_M_ is underestimated even with the highest amount of saturation light (Loriaux et al., 2013). In addition, saturating pulses of ∼2000 µmol photons m^-2^ s^-1^, while sufficient for measuring dark adapted F_M_, may be too low to provide a true F’_M_ under high actinic illumination. In our research we have used a method (McClain and Sharkey, 2020) which involves the use of a single multiphase saturation flash of 7000 µmol photons m^-2^ s^-1^ for 0.3 s, with linearly decreasing light intensity by 20% over another 0.5 s; during this period chlorophyll fluorescence values were recorded. With this method, accurate values of F’_M_ were recorded (Loriaux et al., 2013). The operating (i.e., photochemical) efficiency of PSII in light-adapted state of leaves (Φ_PSII_ or ΔF/F’_M_ ratio) and the electron transport rate were also estimated. Particularly, an estimate of electron transport rate was calculated based on Equation 4 (Motohashi and Myouga, 2015)

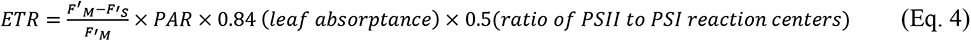

ETR =electron transport rate; F′_M_-maximum, chlorophyll fluorescence; F′_S_=steady state fluorescence; PAR=photosynthetically active radiation at the leaf’s surface. The constant 0.5 corresponds to the excitation energy being divided among both photosystems I and II, and 0.85 is the leaf absorbance coefficient for C4 plants (Oberhuber et al., 1993; Cazzaniga et al., 2013).

### Data Analysis

Analysis of variance was performed to obtain information on the effects of *A. palmeri* sex, light intensity, NPK deficiency and their interactions on the stems (incl. inflorescences) and the leaves; for this, we measured the mineral content, chlorophyll *a* and chlorophyll *b* content, chlorophyll *a*/*b* ratio, as well as chlorophyll fluorescence parameters, namely the steady state fluorescence (F′_S_), maximum fluorescence (F′_M_) whereas the operating efficiency of PSII (Φ_PSII_) and electron transport rate were estimated from these parameters. Before the analysis of stem (incl. inflorescences) and leaf mineral content, mineral concentrations were expressed as a fraction of stem and leaf dry weights (i.e., ppm or µg/g leaf dry weight) to standardize the units of mineral concentrations. The concentration of a mineral in a plant organ can affect the dynamics of another mineral in the same organ (Marschner, 1983). Therefore, an analysis of partial correlations between the leaf and the stem mineral content between *A. palmeri* sex was performed to examine whether the male and the female plants are different in their mineral trade- off strategies. Partial correlation analysis excludes the effects of confounding variables that are numerically related to the variables of interest. In addition, regressions were fitted between chlorophyll *a*, chlorophyll b, chlorophyll *a*/*b* ratio and fluorescence parameters F′_S_, F′_M_ and the estimated Φ_PSII_ against the sampling time throughout the experimental period using white light intensity as a grouping factor. Correlations were also fitted between total chlorophyll and the estimates of Φ_PSII_ and ETR. JMP Pro v. 16.0 (SAS Institute, Cary, NC), and SigmaPlot v. 14.0 (Systat Software, San Jose, CA) were used for statistical analyses and curve-fitting regressions. Means were separated by Least Significant Difference (LSD) test at *a*=0.05 significance level.

## Acknowledgments

We thank Dr. Dimitra A. Loka for insightful comments, Drs. Ananya Sen and Alexendrina (Sandra) Stirbet for suggestions on an earlier draft of this paper.

